# A reported Antarctic environmental microorganism isolated from the blood of sheep on the Mongolian Plateau

**DOI:** 10.1101/2020.07.08.188136

**Authors:** Qianlin Chen, Zheng Cao, Chunyan Li, Maixun Zhu, Shaoqin Zhai, Wengui Fu

## Abstract

Herein, we report a novel *Carnobacterium*-like organism, CS13^T^, isolated from the blood of sheep with persistent diarrhea from a grassland pasturing area in Xilingol League, Inner Mongolia Municipality, China. Homology analysis indicated that CS13^T^ belongs to the genus *Carnobacterium* and is 100% related to the reported environmental microorganism *Carnobacterium antarticum sp. CP1* (*C. CP1*), which was isolated from sandy soil near Davis Station, Antarctica; the following strains are closely related: *Carnobacterium mobile DSM 4848* (97%) and *Carnobacterium funditum DSM 5970* (96%). Similar to those of the *C. CP1*, the short rod-shaped cells of CS13^T^ are 0.4-0.8 μm wide and 1.0-1.5 μm long; exist singly, paired or catenoid; are gram positive, non-spore forming, and facultatively anaerobic; and produce hemolysin. CS13^T^ cannot produce gas or H_2_S but can ferment sucrose, galactose, salicin, and esculin to produce acid. However, in contrast to *C. CP1*, CS13^T^ can produce acid from cellobiose and maltose and is weakly positive for D-mannose fermentation; the growth temperatures range from 20-37°C, the pH range is 5.0-9.0, and the G+C content is 37.84% (4-36°C, pH 6.0-9.5, and 38.1% for *C. CP1*). Furthermore, based on gene annotation analysis, we found that CS13T has 31 more specific genes than *C. CP1* (133 to 102) and that the nonredundant protein similarity to *C. CP1* is only 84.2%. Based on the physiological-biochemical and genetic analysis results, we infer that the organisms isolated from the Mongolian Plateau and sandy soil in Antarctica belong to the same novel species of the genus *Carnobacterium*; therefore, this novel species probably has distributed globally and should not be called *species antarticum*.

## Introduction

Carnobacteria are ubiquitous lactic acid bacteria (LAB), tolerant to freezing/thawing and high pressure and able to grow at low temperatures [1]. The genus belongs to the *family Carnobacteriaceae* of the *phylum Firmicutes*, *class Bacilli*, *order Lactobacillales*, as described in *Bergey’s Manual of Systematic Bacteriology* [2], and includes motile, psychrotolerant, short rod-shaped, gram-positive, facultatively anaerobic, heterofermentative lactic acid bacteria that can produce L-lactic acid from mostly fermented D-glucose [3]. At the time of writing, 12 recognized species had been correctly named and collected in the List of Prokaryotic names with Standing in Nomenclature (LPSN) collection (http://www.bacterio.net).

The species *C. divergens*, *C. gallinarum* and *C. mobile* are frequently encountered in the environment and in foods. *C. antarcticum*, *C. alterfunditum*, *C. funditum* and *C. iners* were isolated from sandy soil, anoxic lake water and the littoral zone of Antarctica [2,4,5]. *C. inhibens* and *C. maltaromaticum* were found in Atlantic salmon and infected Lake Whitefish, respectively [6,7]. Additionally, *C. pleistocenium* was isolated from permafrost of the Fox Tunnel in Alaska [8], *C. viridians was* isolated from vacuum-packed bologna sausage [9], and *C. jeotgali* was isolated from a Korean traditional fermented food [10]. Although a large number of research studies have reported isolation of these bacteria from various regions and environments, many species have not yet been allocated to known species, such as the *Carnobacterium*-like organisms isolated from the larval midgut of a moth species [11], spent mushroom compost [12] and watershed polluted with horse manure [13].

In this study, we isolated a novel *Carnobacterium*-like organism, designated CS13^T^, from the blood of sheep with persistent diarrhea in the Mongolian Plateau in China. To further clarify the diversity of this novel isolated strain and *Carnobacterium antarticum sp. CP1*, this paper discussed the similarities and differences through culture characteristics, phenotypic characterization, and physiological-biochemical and phylogenetic characteristics.

## Materials and methods

### Ethics statement

This study was approved by the Animal Ethics Committee of Chongqing Academy of Animal Sciences. The protocol of blood sample collection was established according to A Good Practice Guide to the Administration of Substances and Removal of Blood, Including Routes and Volumes [14].

### Collection of blood samples and isolation of strains

Grazing sheep with persistent diarrhea were found at Zhenglan Banner, Xilinguole League, Inner Mongolia Municipality, China (N42°42′09″, E116°13′19″), in 2019. Cervical vein blood samples of sheep were collected by a sterile syringe, and the samples were stored in anticoagulative tubes at 4°C. To culture and separate the pathogens from blood samples, brain heart infusion broth liquid (BHI) medium and Columbia blood agar (CBA, supplemented with 5% (v/v) defibrinated sheep blood) medium were prepared as previously described [15]. Aliquots of 100 μL of the blood samples were streak-inoculated on CBA medium at 4°C, 20°C, 25°C, 30°C or 37°C in the presence or absence of oxygen. Colonies were observed after 72 h, and the clearest colonies were subcultured into BHI medium and then cultured for 48 h. Recovered pure cultures were preserved at −80°C in BHI broth supplemented with 20% glycerol.

### Physiology and biochemistry observation

To define the optimal culture conditions, the isolated strain was inoculated in BHI medium with extra NaCl concentrations of 1.0-10.0% (at intervals of 1.0%, w/v) at pH 5.0-10.0 (at intervals of 1.0) at growth temperature for 72 h separately. CBA was used as a growth and hemolysin examination medium to culture the isolated strain. Gram staining was conducted with a gram staining kit (Solarbio) and observed by optical microscopy (Nikon). The morphology, size and flagellum ultrastructure of the isolated strain were observed by a JEOL JEM-1200EX electron microscope after uranyl acetate and citromalic acid lead double staining. The biochemical properties, including glycolysis reaction, indole production, hydrogen sulfide production, methyl red test, pyruvate utilization, nitrate reduction and acid production, were determined using a Micro-Biochemical Identification Tube (Hopebio).

### Homology and phylogenetic analyses

The genomic DNA of the isolated strain was extracted using a Bacterial DNA Kit (TIANGEN) and then submitted to Sangon Biotech (Shanghai) for sequencing. Homologous sequences were compared with NT (NCBI nucleotide sequences database), NR (NCBI nonredundant protein sequences database) and Swiss-Prot (manually annotated and reviewed protein sequences database). Phylogenetic analysis was performed via maximum-likelihood, maximum-parsimony and neighbor-joining algorithms in MEGA version 7.0 [16]. Additionally, comparisons of the core genes, dispensable genes and specific genes were also used in phylogenetic analyses.

## Results

### Isolation and identification

Earlier isolates on CBA medium incubated with oxygen at 20-37°C (optimum, 30°C) for 72 h presented bacterial colonies 1-2 mm in diameter that were white-gray and opaque, with neat edges; had a smooth convex elevation and were surrounded by a tiny hemolysis halo. Furthermore, the growth of bacteria was observed at pH 5.0-9.0 (optimum pH=8.0) and in the presence of 0-5% (w/v) NaCl (optimum, 1%) when the isolated strain was inoculated in BHI media with different pH and salinity values. Electron microscopy demonstrated that the cells of the isolated strain were slightly curved short rods approximately 0.4-0.8 μm wide and 1.0-1.5 μm long, with flagella occurring singly or in pairs or short chains (Fig 1).

**Fig 1.**
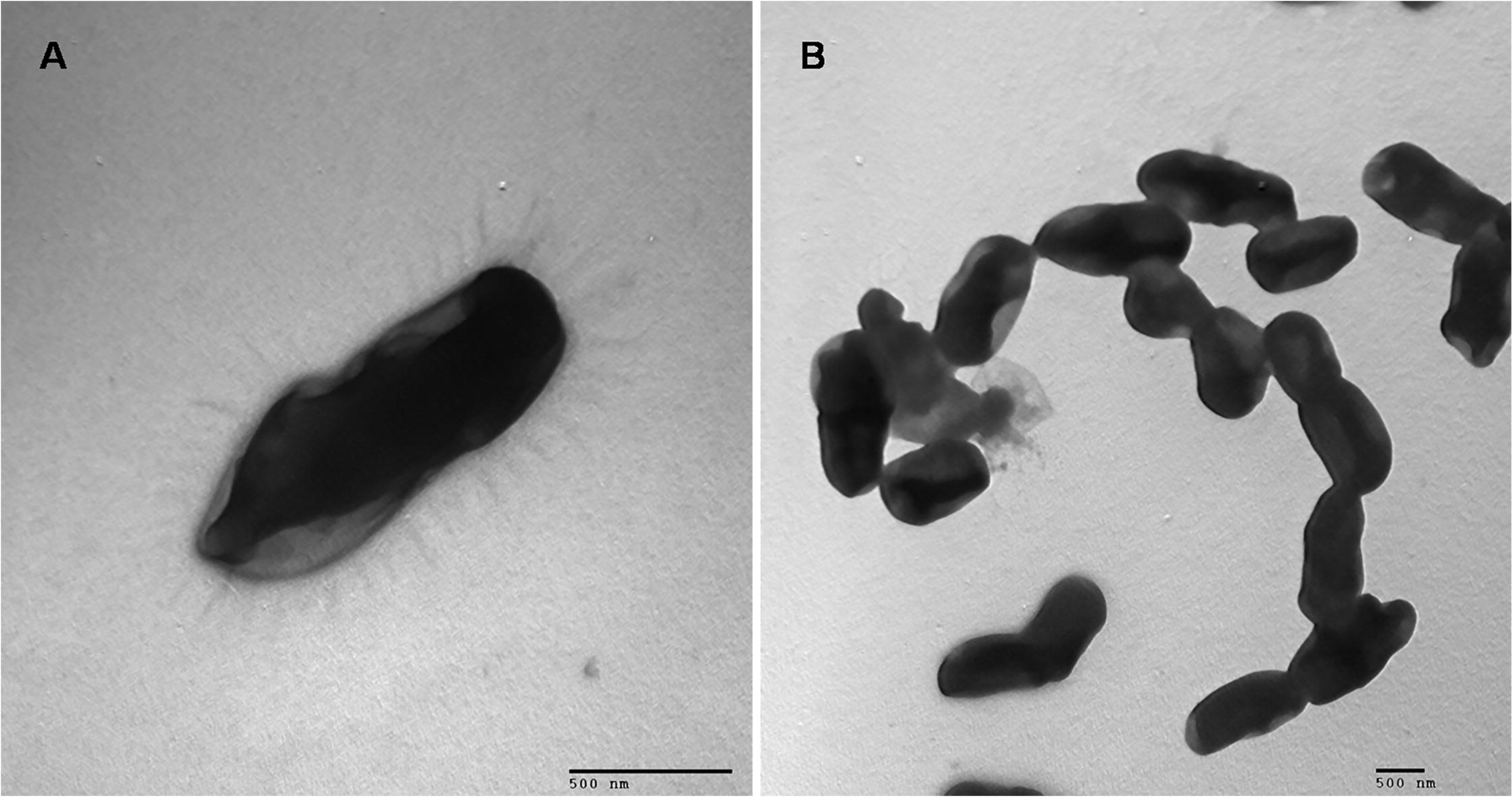
Electron Microscope Observation. Negative stain electron microscopy of single (A) and pairs (B) of isolated strains.

### Physiology and biochemistry

The isolate CS13^T^ has biochemical and physiological characteristics similar to those of the Antarctica-isolated strains *Carnobacterium antarticum sp. CP1* and *Carnobacterium funditum DSM 5970* and the frozen meat-isolated strain *Carnobacterium mobile DSM 4848*. The four strains exhibit short rod shapes, positive Gram staining, motility, facultatively anaerobic growth, growth at low temperatures, negative oxidase and catalase activities, and no H_2_S production. In contrast, *C. DSM 4848* is the only strain that produces gas, and except *C. DSM 5790*, all of them utilized esculin, D-glucose, D-mannose, N-acetyl-glucosamine, salicin and sucrose to produce acid. The physiological similarity between the isolates and their closest relatives among the genus are presented in Table 1.

**Table 1.**
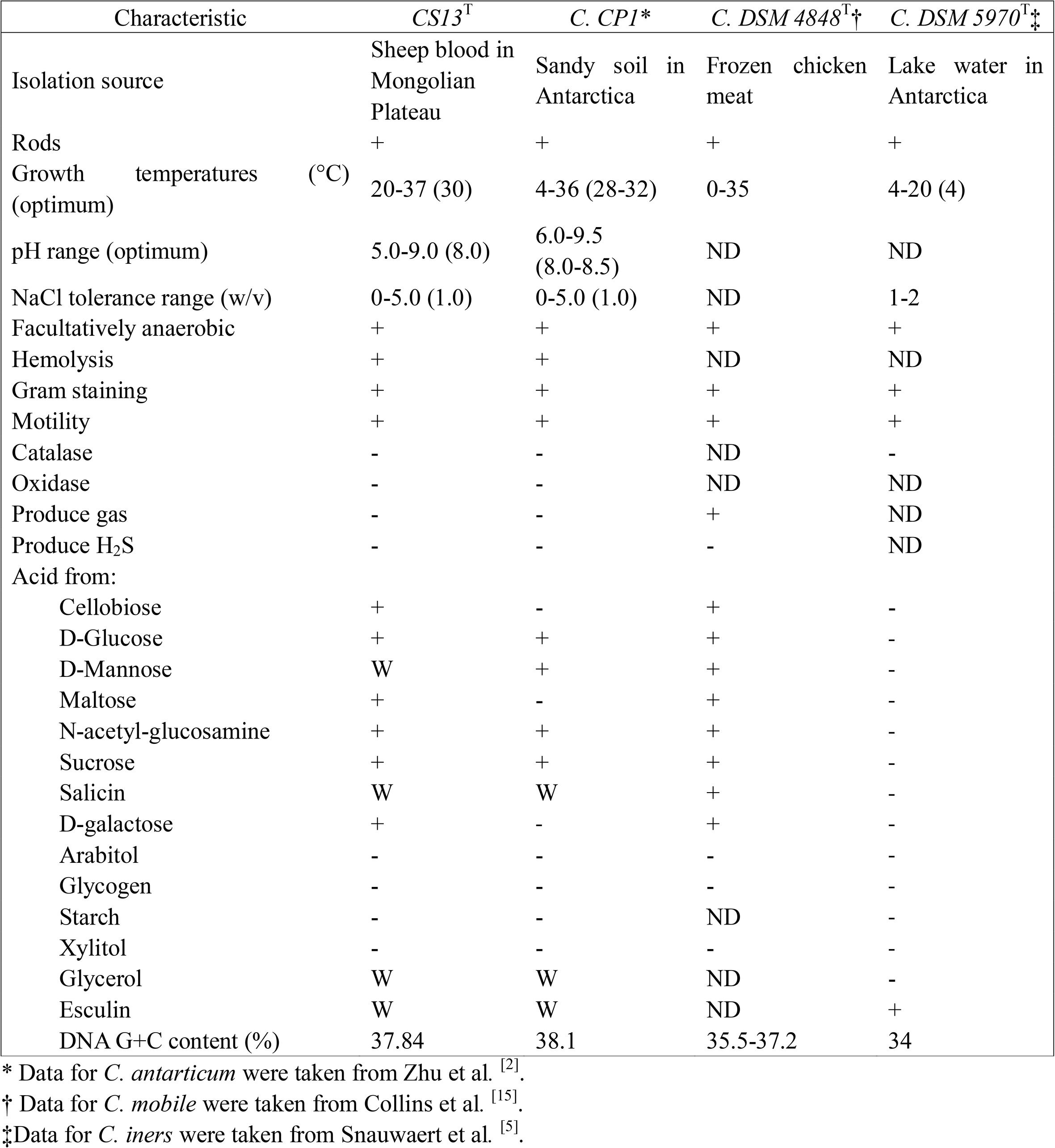
Differential Physiological Characteristics Distinguishing Strain CS13^T^ from the Closest Members of the Genus *Carnobacterium.* +, −, W and ND Represent Positive, Negative, Weakly Positive and ‘No Data’, Respectively.

### Homology analysis

The 16S rRNA gene of CS13^T^ was sequenced by Sangon Biotech (Shanghai). 16S rRNA sequence alignment (NCBI blastn) showed that CS13^T^ shares 100% identity with *Carnobacterium antarticum sp. CP1*, 97% with *Carnobacterium mobile DSM 4848* and 96% with *Carnobacterium funditum DSM 5970.* The 16S rRNA gene and pangenome phylogenetic trees (Fig 2A and B) also demonstrated that CS13^T^ and CP1 have the closest relationship. Therefore, we infer that the organism isolated from the blood of sheep on the Mongolian Plateau belongs to the genus *Carnobacterium* and that CS13^T^ is on the same branch as *C. CP1*.

**Fig 2.**
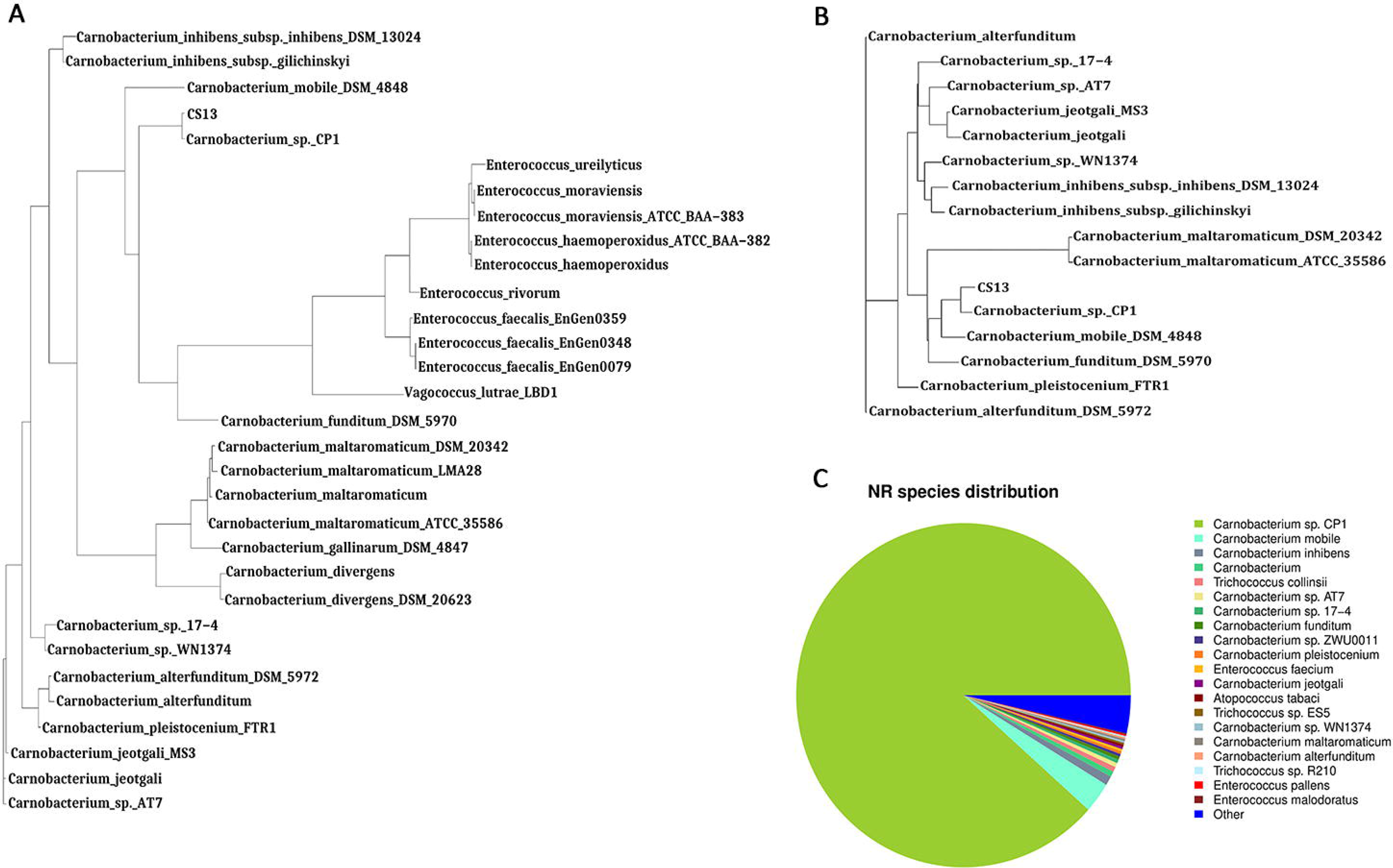
Homology Analysis. Evolutionary relationships based on the 16S rRNA Gene (A) and pangenome (B) of strain CS13^T^ and related species. (C) The nonredundant protein gene annotation of strain CS13^T^.

In contrast, the NR species distribution results indicated that CS13^T^ had only 84.2% of protein coding genes that matched those of *C. CP1* (Fig 2C). Orthologous cluster analysis revealed that CS13^T^ has 133 specific genes, in comparison with *C. CP1* (102 specific genes) and other related species (Fig 3A and B), indicating that strain CS13^T^ could be classified as a novel species of the genus *Carnobacterium*.

**Fig 3.**
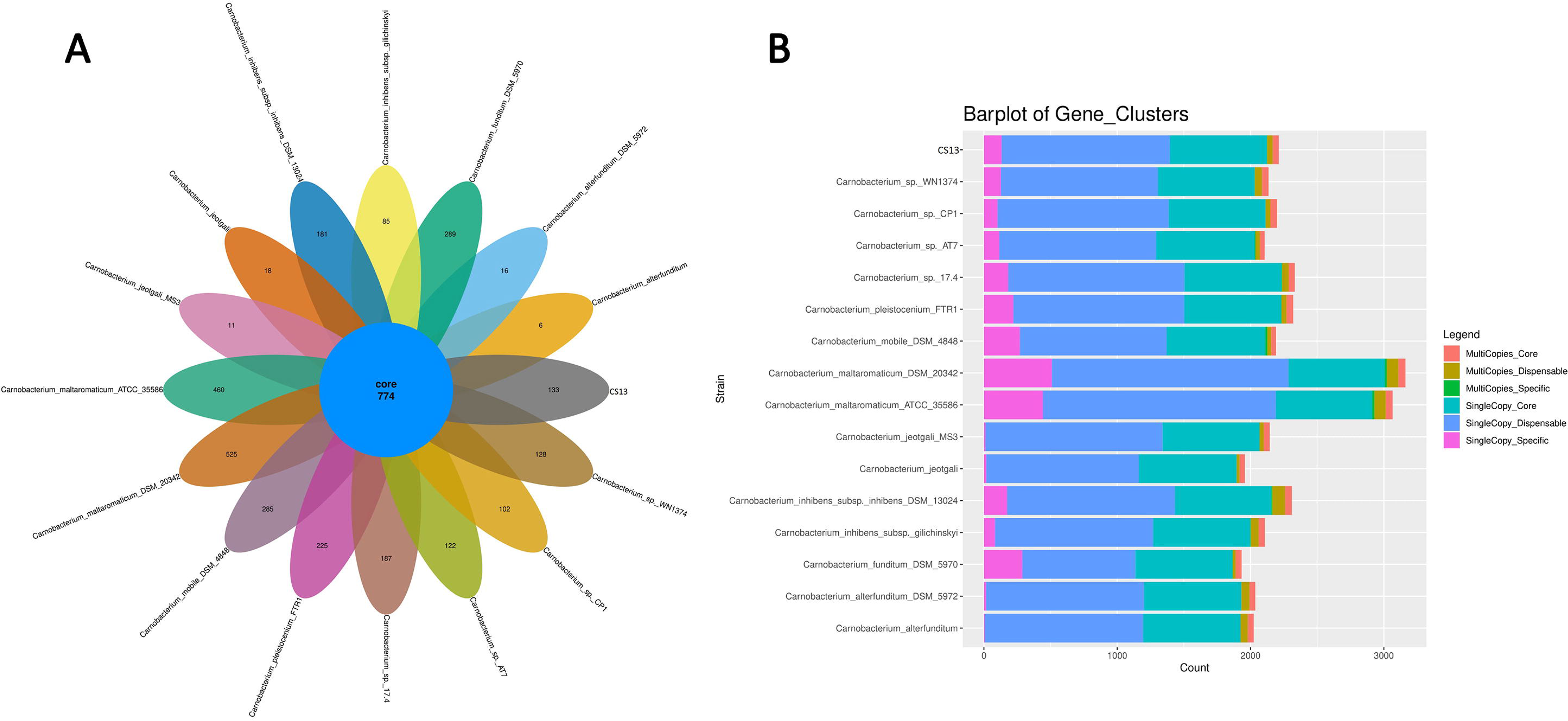
Pan Genomic Analysis. (A). Quantity of core genes (center) and specific genes (petal) of strain CS13^T^ and related species. (B). Gene clusters of strain CS13^T^ and related species.

The protein sequence alignment of CS13^T^ and *C. CP1* was carried out by the Circos package (Fig 4). The results illustrated that although the protein sequences of CS13^T^ and *C. CP1* are also highly homologous, there are many specific fragments of CS13^T^, such as NODE_88, NODE_101, NODE_113, NODE_115 and NODE_145, which further confirms CS13^T^ as a novel species.

**Fig 4.**
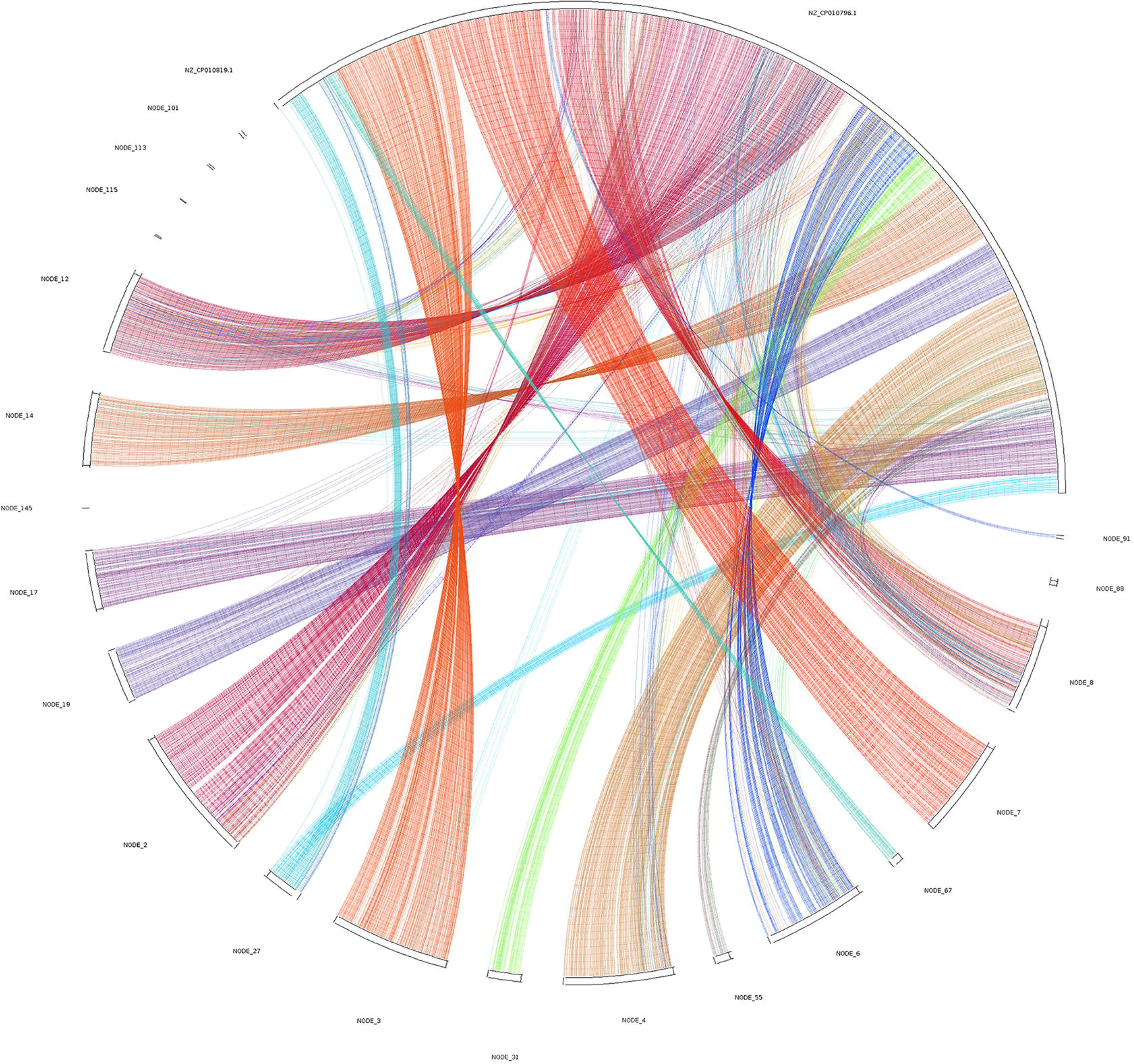
Circos Package. The protein sequence alignment of CS13^T^ and *C. CP1.*

## Discussion

*Carnobacterium* is found mostly in the intestines of animals. As intestinal probiotics, certain *Carnobacterium* can also effectively inhibit pathogens and spoilage microorganisms, so they are widely used as a food additive. Among 12 species of *Carnobacterium*, *C. divergens* and *C. maltaromaticum* are frequently isolated from food, particularly in vacuum-packaged (VP) meat and meat products, which also show the ability to inhibit pathogenic and spoilage microorganisms in diverse food matrices. Other scholars reported that *C. maltaromaticum* B26 and *C. divergens* B33, isolated from the intestine of healthy rainbow trout, are beneficial for enhancing cellular and humoral immune responses, and they were selected as potential probiotics with effectiveness. In this study, we first isolated a *Carnobacterium antarticum sp. CP1-like* organism, CS13^T^, from the blood of sheep with persistent diarrhea in the Mongolian Plateau in China. The results of the physiological-biochemical and genetic analyses indicate that the organisms isolated from the Mongolian Plateau and Antarctica belong to the same novel species of the genus *Carnobacterium*. However, preliminary results prove that this novel species should not be called *species antarticum* and suggest that this species is probably distributed globally and adapts to varying areas and surroundings. Whether this novel species has bacteriostatic effects or can be used in the food industry or other fields remain to be further studied.

## References

1. Leisner JJ, Laursen BG, Prévost H, Drider D, Dalgaard P. Carnobacterium: positive and negative effects in the environment and in foods. FEMS Microbiol Rev. 2007;31: 592–613.

2. Zhu S, Lin D, Xiong S, Wang X, Xue Z, Dong B, et al. Carnobacterium antarcticum sp. nov., a psychrotolerant, alkaliphilic bacterium isolated from sandy soil in Antarctica. Int J Syst Evol Microbiol. 2018;68: 1672–1677.

3. Nicholson WL, Zhalnina K, de Oliveira RR, Triplett EW. Proposal to rename *Carnobacterium inhibens* as *Carnobacterium inhibens* subsp. inhibens subsp. nov. and description of *Carnobacterium inhibens* subsp. gilichinskyi subsp. nov., a psychrotolerant bacterium isolated from Siberian permafrost. Int J Syst Evol Microbiol. 2015;65: 556–561.

4. Franzmann PD, Hpfl P, Weiss N, Tindall BJ. Psychrotrophic, lactic acid-producing bacteria from anoxic waters in Ace Lake, Antarctica; *Carnobacterium funditum* sp. nov. and Carnobacterium alterfunditum sp. nov. Arch Microbiol. 1991;156: 255–262.

5. Snauwaert I, Hoste B, De Bruyne K, Peeters K, De Vuyst L, Willems A, et al. Carnobacterium iners sp. nov., a psychrophilic, lactic acid-producing bacterium from the littoral zone of an Antarctic pond. Int J Syst Evol Microbiol. 2013;63: 1370–1375.

6. Cantas L, Fraser TWK, Fjelldal PG, Mayer I, Sørum H. The culturable intestinal microbiota of triploid and diploid juvenile Atlantic salmon (Salmo salar) - a comparison of composition and drug resistance. BMC Vet Res. 2011;7: 71.

7. Loch TP, Xu W, Fitzgerald SM, Faisal M. Isolation of a *Carnobacterium maltaromaticum*-like bacterium from systemically infected lake whitefish (*Coregonus clupeaformis*). FEMS Microbiol Lett. 2008;288: 76–84.

8. Pikuta EV, Marsic D, Bej A, Tang J, Krader P, Hoover RB. Carnobacterium pleistocenium sp. nov., a novel psychrotolerant, facultative anaerobe isolated from permafrost of the Fox Tunnel in Alaska. Int J Syst Evol Microbiol. 2005;55: 473–478.

9. Holley RA. Carnobacterium viridans sp. nov., an alkaliphilic, facultative anaerobe isolated from refrigerated, vacuum-packed bologna sausage. Int J Syst Evol Microbiol. 2002;52: 1881–1885.

10. Kim MS, Roh SW, Nam YD, Yoon JH, Bae JW. Carnobacterium jeotgali sp. nov., isolated from a Korean traditional fermented food. Int J Syst Evol Microbiol. 2009;59: 3168–3171.

11. Shannon AL, Attwood G, Hopcroft DH, Christeller JT. Characterization of lactic acid bacteria in the larval midgut of the keratinophagous lepidopteran, Hofmannophila pseudospretella. Lett Appl Microbiol. 2001;32: 36–41.

12. Ntougias S, Zervakis GI, Kavroulakis N, Ehaliotis C, Ehaliotis C, Papadopoulou KK. Bacterial diversity in spent mushroom compost assessed by amplified rDNA restriction analysis and sequencing of cultivated isolates. Syst Appl Microbiol. 2004;27: 746–754.

13. Simpson JM, Santo Domingo JW, Reasoner DJ. Assessment of equine fecal contamination: the search for alternative bacterial source-tracking targets. FEMS Microbiol Ecol. 2004;47: 65–75.

14. Diehl KH, Hull R, Morton D, Pfister R, Rabemampianina Y, Smith D, et al. A good practice guide to the administration of substances and removal of blood, including routes and volumes. J Appl Toxicol. 2001;21: 15–23.

15. Zhu S, Wang X, Zhang D, Jing X, Zhang N, Yang J, et al. Complete genome sequence of hemolysin-containing *Carnobacterium* sp. strain CP1 isolated from the Antarctic. Genome Announc. 2016;4: e00690–16.

16. Collins MD, Farrow JAE, Phillips BA, Feresu S, Jones D. Classification of Lactobacillus divergens, Lactobacillus piscicola, and some catalase-negative, asporogenous, rod-shaped bacteria from poultry in a new genus, Carnobacterium. Int J Syst Evol Microbiol. 2008;58: 2672–2672.

